# hnRNP A1 induces aberrant CFTR exon 9 splicing *via* a newly discovered Exonic Splicing Silencer element

**DOI:** 10.1101/2023.08.24.554598

**Authors:** Christelle Beaumont, Ming-Yuan Chou, Cristiana Stuani, Emanuele Buratti, Peter Josef Lukavsky

**Affiliations:** CEITEC-Central European Institute of Technology, Masaryk University, Kamenice 5, 625 00 Brno, Czech Republic; International Centre for Genetic Engineering and Biotechnology, Padriciano 99, 34012 Trieste, Italy

**Author notes:** (C.B.); (M.Y.C.); (P.J.L.). (C.S.); (E.B.).

**Keywords:** Alternative splicing, CFTR, pre-mRNA, RNA binding protein, RNA-binding mode, RRM (RNA Recognition motif), NMR spectroscopy

## Abstract

RNA-protein interactions play a key role in the aberrant splicing of CFTR exon 9. Exon 9 skipping leads to the production of a non-functional chloride channel associated with severe forms of cystic fibrosis. The missplicing depends primarily on variations in the polymorphic (TG)_m_T_n_ locus upstream of exon 9. At the pre-mRNA level, it generates an extended UG-rich binding site for TDP-43, associated with hnRNP A1 recruitment, and prevention of exon 9 3’ splicing site (3’ss) recognition. While TDP-43 is the dominant inhibitor of exon 9 inclusion, the role of hnRNP A1, a protein with two RNA recognition motifs (RRM1 and RRM2) and a glycine-rich domain, remained unclear. In this work, we have studied the interaction between hnRNP A1 and the CFTR pre-mRNA using NMR spectroscopy and Isothermal Thermal Calorimetry (ITC). The affinities are submicromolar and ITC data suggest that the separate RRMs as well as tandem RRMs form 1:1 complexes. NMR titrations reveal that hnRNP A1 interacts with model CTFR 3’ss sequences in a fast exchange regime at the NMR timescale. Splicing assays finally show that this hnRNP A1 binding site represents a previously unknown exonic splicing silencer element. Together, our results shed light on the mechanism of aberrant CFTR exon 9 splicing.

## 1. Introduction

Cystic fibrosis is an autosomal recessive, inherited disease that affects the respiratory, digestive, and reproductive systems^1^. The CFTR (Cystic Fibrosis Transmembrane conductance Regulator) gene encodes for the CFTR protein which acts as a chloride channel in epithelial cell membranes^2^. The CFTR gene contains 27 exons. All exons included in the mRNA produces a functional protein^3-5^. The expression of most eukaryotic genes requires mRNA splicing which removes introns and ligates exons to produce a mature mRNA. Two types of regulating, cis-acting elements are found in pre-mRNAs which either enhance or silence splicing *via* interaction with specific trans-acting splicing factors. The cis-regulatory elements located upstream or downstream of the regulated exon within the pre-mRNA, define which exons are included or excluded from the pre-mRNA^6^. Silencer elements interact with negative trans-acting factors such as hnRNPs (heterogeneous nuclear ribonucleoproteins) and thus repress splicing. The hnRNP family is conserved in humans which highlights their importance for functioning in pre-mRNA maturation^7^.

In aberrant splicing of the CFTR gene, there is limited inclusion of exon 9 in the final mRNA due to a polymorphic sequence in the intron upstream of this exon. The result is a defect in the nucleotide binding domain of the CFTR channel which leads to CFTR channel dysfunction. Upstream of exon 9, in intron 8, polymorphisms can produce different variant numbers of the (TG)_m_(T)_n_ element^8-9^. At the pre-mRNA level, it has been observed that polymorphic (UG)_m_(U)_n_ repeats with low number of Us and concomitant high number of UG repeats can repress the use of the polypyrimidine tract by altering the interaction with trans-acting proteins.

In regular splicing, the small U1 snRNA of the snRNP interacts with the 5’ss of the pre-mRNA and is supported by auxiliary factors such as the serine-arginine-rich proteins (Figure 1A). Recognition of the 3’ss is conducted by a concerted interaction of the splicing factor 1, also called branch point binding protein (SF1/BBP) to the branch point sequence and the heterodimeric auxiliary factor U2AF. While UA2F35 recognizes the conserved AG dinucleotide at the 3’ss, U2AF65 binds the polypyrimidine tract located upstream^10-13^.

**Figure 1.**
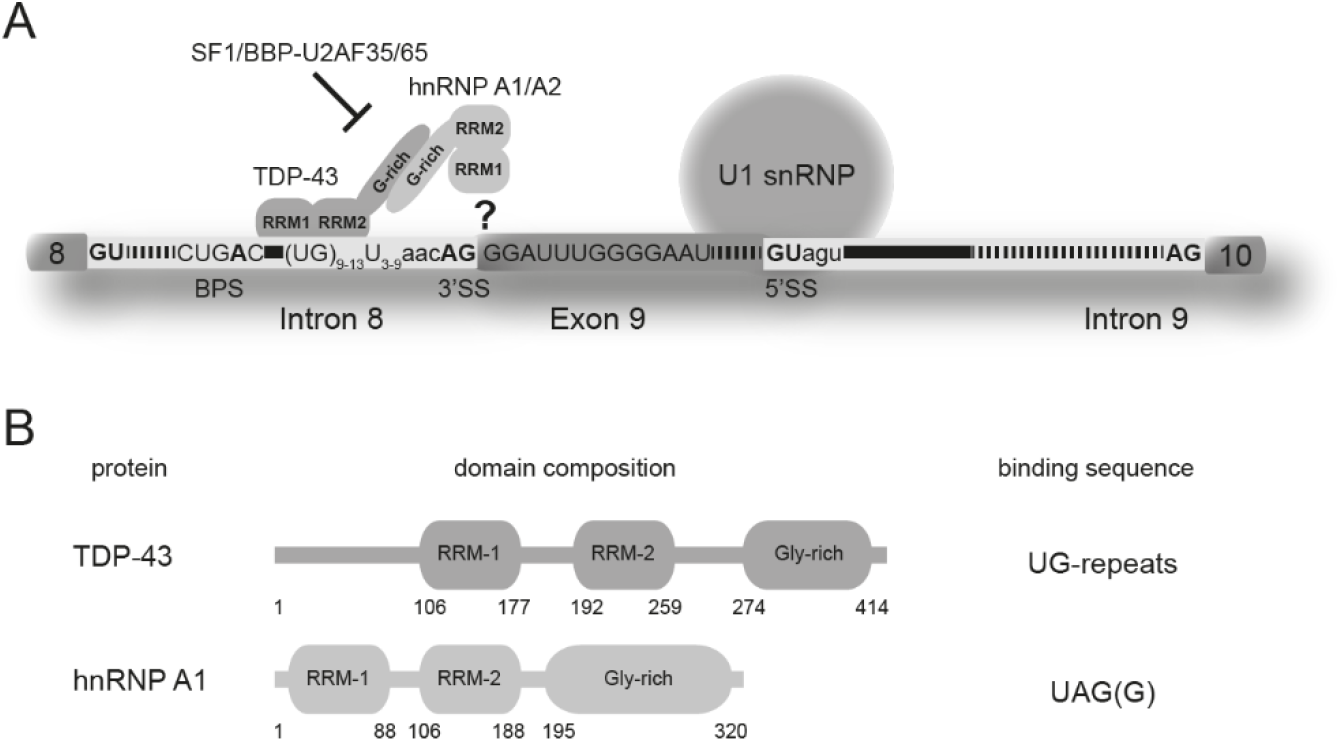
Aberrant splicing of CFTR exon 9. (**A**) Recognition of the CFTR 3’ss by TDP-43 and hnRNP A1. The relevant RNA sequences in intron 8 and exon 9 are displayed. TDP-43 binds the elongated UG-repeat and recruits hnRNP A1 via their C-terminal glycine-rich domains. Access of SF1/BBP and U2AF35/65 to the canonical binding sites (BPS, polypyrimidine tract and 3’ss AG) at the 3’ss are impaired. The pre-mRNA interaction site of hnRNP A1 is unknown. (**B**) Schematic domain representation of human TDP-43 RBD and human hnRNP A1. Domain boundaries of each domain are indicated according to the full length protein sequences (NP_031401 and NP_112420.1).

Studies showed that TDP-43 protein, displaying two RRM domains and an unstructured, glycine-rich C-terminus, has a high affinity for a (TG)_12_T_5_ sequence and binds to this site at the 3’ss of CFTR exon 9^14-15^. Since this polymorphism is associated with a very short polypyrimidine tract, binding of the canonical factor U2AF65/35 is presumably impaired (Figure 1A). TDP-43, on the other hand, has a strong affinity for the concomitant, extended UG repeat and thus acts as the principal inhibitor causing aberrant splicing of CFTR exon 9^16-17^.

Previous studies have shown that beside the RRM domains, the C-terminal of TDP-43 protein is also essential for aberrant CFTR exon 9 splicing *via* an interaction with the C-terminus of hnRNP A1 (Figure 1A-B)^18^. Using deletion mutants, Baralle, Buratti and colleagues, mapped the minimal region (region between 321-366 of the TDP-43 C-terminus) required for the interaction of hnRNP A1 and TDP-43^19^. Interestingly, similarly to TDP-43 the hnRNP A1 protein also contains two RRM domains and an unstructured, glycine-rich C-terminus (Figure 1B). The latter provides a platform for protein-protein interactions playing a key role in regulating gene expression including alternative splicing, nuclear and export from the nucleus to the cytoplasm, telomere maintenance, mRNA stability and turnover, mRNA processing and translation regulation^20^. Although TDP-43 has been shown to be the main splicing inhibitor, hnRNP A1 is also required for aberrant CFTR exon 9 splicing^18^. However, until present the hnRNP A1 interaction site of its tandem RRMs at the 3’ss of exon 9 of the CFTR pre-mRNA remained elusive (Figure 1A).

The tandem RRMs of human hnRNP A1 are connected by a 17 amino acid linker and provide a platform for RNA interactions (Figure 1B). The tandem RRMs have been crystallized in the free and bound form where they dimerize upon binding with ssDNA. Two ssDNA sequences bind to each tandem RRM in an antiparallel manner where the 5’ extremity of the ssDNA binds to the RRM1 of one tandem RRM domain while the 3’ extremity binds to the RRM2 of the second copy^21^. The presence of this specific dimer arrangement and binding mode might be due to the packaging forces in the crystals. A study in solution using NMR spectroscopy combined with segmental isotope labeling shows that the orientation of the unbound tandem RRMs differs from the crystal structure of the free form and resembles more the one found in the ssDNA-bound form^22^. However, the interaction between the tandem RRMs and RNA remained controversial. Despite all the structural insights from model RNA-hnRNP A1 complexes, no interaction with a naturally occuring splice site has been characterized in detail.

Our study set out to decipher the role of hnRNP A1 in aberrant splicing of CFTR exon 9. Splicing assays reveal a previously unrecognized ESS within CFTR exon 9. NMR titration experiments and ITC measurements confirm that this element interacts with hnRNP A1 with submicromolar affinity in a specific manner forming 1:1 complexes. Our results explain how TDP-43 and hnRNP A1 work in concert to block formation of a splicing competent complex at the CFTR exon 9 3’ss and thus cause o aberrant CFTR exon 9 splicing.

## 2. Materials and Methods

### 2.1. Cloning, expression and purification of hnRNP A1 tandem RRMs

The sequence of DNA encoding the tandem RRMs composed of 196 amino-acids were subcloned by PCR amplification into the pET28a vector. The construct contains a 2xHis_6_ tag, a lipoyl domain followed by a TEV protease cleavage site for the tag and the lipoyl domain to be removed after the first step of purification. Recombinant proteins were overexpressed in *E. coli* BL21(DE3) codon plus cells (Novagen) in LB rich or M9 minimal media supplemented with ^15^NH_4_Cl or ^15^NH_4_Cl and ^13^C-glucose. The cells were grown at 37°C to OD600 ∼0.6 and then at 20°C for 1h. Protein expression was induced at OD600 ∼0.9 by addition of 1 mM isopropyl-β-D-thiogalactopyranoside (IPTG) and further incubated for 15h at 20°C. Cells were harvested after 15h and then centrifuged at 4000 g for 20 min at 4°C. The cell pellet was resuspended in lysis buffer A (20 mM HEPES pH 7.5, 1 M NaCl, 30 mM imidazole, 5 mM BME, 10% (w/v) glycerol) supplemented with protease inhibitors, and lysed by high pressure with a french press. The cell lysate was centrifuged 30 min at 39,000 g at 4°C. The supernatant was loaded on a Ni-NTA column on a ÄKTA Prime purification system (GE Healthcare), and after washing with buffer A the protein of interest was eluted with 300 mM imidazole gradient. The fractions containing the protein of interest were collected, then TEV protease was added according the estimation of the amount of the protein and then dialyzed against a buffer A without imidazole overnight at room temperature to perform the cleavage of tag. Then the sample was reloaded onto a Ni-NTA column to remove the tag, TEV protease and uncleaved protein. Samples containing the protein were further purified by size exclusion chromatography on a Superdex 75 column equilibrated with NMR buffer (50 mM sodium phosphate pH 6.5, 1mM BME, 50 μM EDTA) at 4°C and concentrated to 1 mM with a Vivaspin 10.000 MWCO.

### 2.2. Cloning, expression and purification of RRM1 and RRM2 of hnRNP A1

The sequence of DNA encoding RRM1 and RRM2 individually (RRM1 from residues 1-105 and RRM2 from residues 91-196) were subcloned by PCR amplification into the same modified pET28a vector as above. Recombinant proteins were overexpressed and purified as for the construct containing both RRMs. Samples containing proteins were further purified by size exclusion chromatography on Superdex 75 column equilibrated with NMR buffer (25 mM sodium phosphate pH 6.5, 1mM BME, 50 uM EDTA) at 4°C and concentrated to 1 mM with a Vivaspin 10.000 MWCO.

### 2.3. ssDNA purification and desalting

The ssDNAs were purchased from Dharmacon. The 15-mer ssDNA (5’-CAGGGAT-TTGGGGAC-3’), the 7-mer ssDNAs (5’-CAGGGAT-3’) and the three 8-mer ssDNAs (5’-TGGGGAAT-3’, 5’-CAGGGATC-3’, 5’-CTGGGCAC-3’) were purified under denaturing conditions using the ultiMate 3000 HPLC system with an anion-exchange preparative DNAPac PA100 Nucleic Acid Column (both Thermo Fisher Scientific) as described before^23^. We performed the purification at 85°C with a flow rate of 20ml/min. The column was equilibrated with the buffer containing 6M urea, 12.5 mM Tris/HCL, pH 7.4. The sample was loaded into the sample loop and then eluted from the column with the buffer containing 6M urea, 12.5 mM Tris/HCL, 500 mM NaClO4, pH 7.4 up to 50% of NaClO4 while collecting 10m ml fractions. The fractions were analyzed by SDS-PAGE (20% acrylamide, 8M urea). The ssDNA was visualized with 0.1% toluidine. The fractions containing the ssDNA were desalting using a ÄKTA prime FPLC system (GE Healthcare) equipped with three 5 ml DEAE weak anion-exchange columns in series^24^. The columns were equilibrated with the buffer containing 20 mM ammonium bicarbonate, pH 7.5 (degassed for 30 min). The sample was diluted twice, then loaded into the sample loop and injected with the buffer at a flow rate of 5 mL/min. A steep gradient containing 100% of 2.53 M ammonium bicarbonate, pH 8 was applied to collect 10 mL fractions. Fractions with the ssDNA were collected and then lyophilized.

### 2.4. NMR experiments

All NMR experiments were carried out using Bruker Avance III HD 700, 850 and 950 MHz spectrometers each equipped with cryoprobes. Data acquisition was performed at 298K, samples were measured in NMR buffer with 10% D_2_O. Data were processed using Topspin 3.2*/*3.5 (Bruker) and analyzed with Sparky (http://www.cgl.ucsf.edu/home/sparky/). Protein resonance assignments were taken from BioMagResBank under accession number 18728^22^. The buffer used (25 mM sodium phosphate pH 6.5, 1mM DTT) was similar to our NMR buffer (see above) so that resonance assignments could be directly transferred to our spectra. CSPs from NMR titration experiments (2D ^1^H-^15^N HSQC spectra) were calculated using the formula: Δδ=[(δH^N^)^2^ + (δN/6.51)^2^]^1/2^. HSQC titrations were carried out using ^15^N-labeled hnRNP A1 protein constructs and unlabelled ssDNA. Spectra were recorded at the following molar ratios Protein/ssDNA: 1/0.2; 1/0.4; 1/0.6; 1/0.8; 1/1; 1/2.

### 2.5. ITC measurements

ITC experiments were performed on a VP-ITC instrument (Microcal) at 25°C. The calorimeter was calibrated according to the manufacturer’s instructions. Protein and ssDNA samples were dialyzed against the NMR buffer. Concentrations of proteins and ssDNAs were determined using optical absorbance at 280 and 260 nm, respectively. The ssDNA (200 μM) was injected into the sample cell containing the protein at a concentration of around 20 μM. Titrations consisted of 20 injections of 2 uL except for the first injection (1.5 uL) with a 2 min spacing between each injection. Data were fitted with one site binding model using Microcal Origin.

### 2.6. Splicing assay

Plasmid TG11-T5 has been previously described by Niksic et al.^25^. Mutations were inserted using complementary nucleotides and Liposome-mediated transfections of 3x10^5^ human HeLa cells were performed using DOTAP Liposomal Transfection Reagent (Alexis Biochemicals) according to manufacturer instructions. After 18 hours the transfectiom medium was replaced with fresh medium and 24 hours later the cells were washed with PBS and RNA was purified using RNAwiz (Ambion). RT-PCR reactions to specifically amplify the minigene transcripts was performed as previously reported^26^.

## 3. Results

### 3.1. In vivo splicing assay identifies a novel ESS in CFTR exon

The hnRNP A1 protein binds the SELEX sequence 5’-UAGGGA/U-3’^27^ with high affinity and iCLIP studies of functional hnRNP A1 binding sites further indicate specificity for a 5′-UAGG-3′ motif^28^. Inspection of the CFTR 3’ss (5’-AAC**<u>AG</u>GGAU**UU**GGGGAAU**-3’) displays similar AG-rich motifs at the conserved 3’ss **<u>AG</u>** and within the first codons of exon 9 (Figure 1A).

To test the effect on splicing of these motifs, we altered the RNA sequence in exon 9 while keeping its coding potential for amino acids. The universal 3’ss **<u>AG</u>** was not mutated since it would delete the 3’ss but the second motif was altered progressively to delete the AG-rich motif (Figure 1A). The mutations were introduced into a CFTR exon 9 splicing minigene reporter system which contains the exon 9 sequence, the splicing junctions and part of the flanking introns with the TG11-T5 repeat at the 3’ss constituting the functional TDP-43 binding site which is indispensible for aberrant CFTR exon 9 splicing as previously described (Figure 2A)^15, 19, 29^. In contrast to previous studies, the splicing assay was not coupled with RNAi-mediated knockdown of endogenous TDP-43 or hnRNP A1 and add-back of siRNA-resistant wild-type (wt) proteins so that the effect on splicing could be studied in a natural protein background. In addition, it was already previously shown that overexpressing hnRNP A1 in a TG11-T5 minigene context was able to increase the level of exon 9 skipping^26^ and that the knockdown of hnRNP A1 would lead to 100% exon inclusion since both TDP-43 and hnRNP A1 are required for exon 9 skipping^18-19^. As a last consideration, we opted for the TG11-T5 repeat minigene because it reflects a borderline physiological polymorphic situation in humans. Minigenes with higher TG repeats (i.e. TG13) were not considered since the resulting increased binding of TDP-43 to these repeats could have masked the inhibitory role of hnRNP A1.

**Figure 2.**
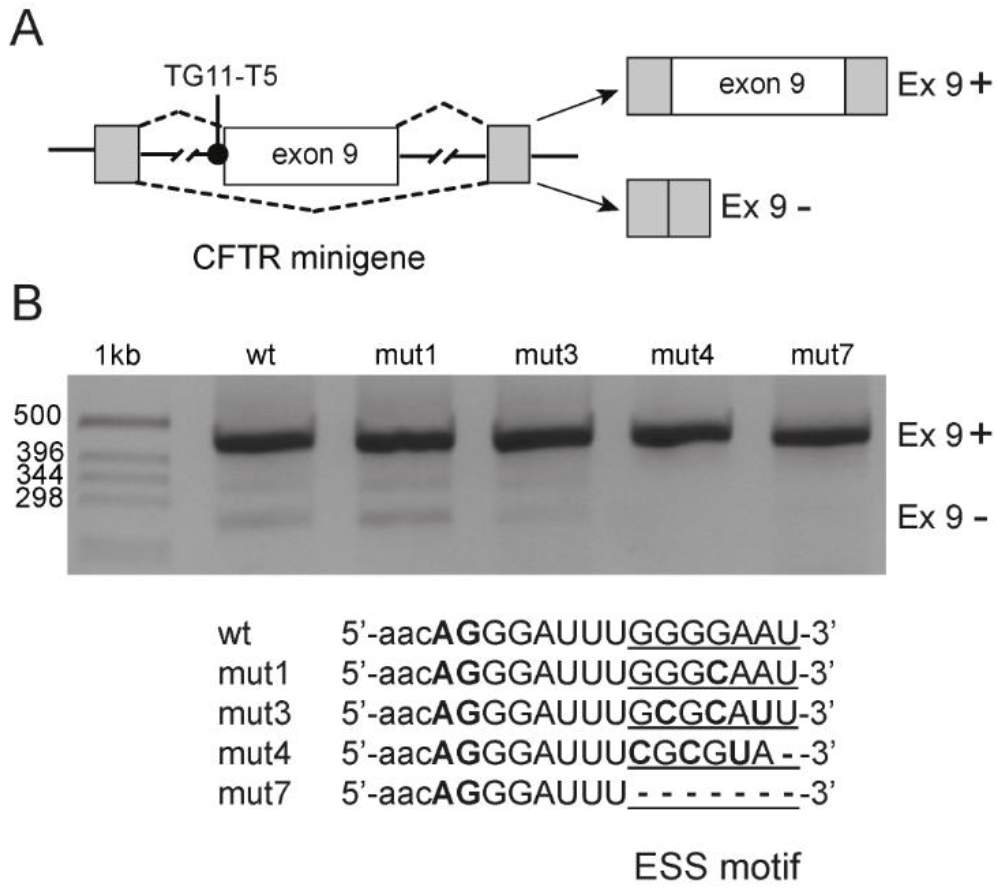
Minigene splicing assay identifies a novel ESS motif in CFTR exon 9. (**A**) Schematic representation of the CFTR exon 9 minigene construct used in the transfection experiments. Lines represent introns and boxes represent exons. The intronic TG11-T5 TDP-43 binding site upstream of exon 9 is also indicated. Dashed lines indicate alternative pre-mRNA processing of the minigene and the two major splice forms are shown on the right. (**B**) CFTR exon 9 inclusion upon mutation of the ESS motif in exon 9. The TG11-T5 CFTR exon 9 reporter minigenes (wt and mutants mut1, mut3, mut4, mut7) were transfected in HeLa cells. Progressive mutations of the ESS motif lead to full inclusion of exon 9.

The wt 5’-GGGGAAU-3’ motif and the mutant Gly to Gln codon change (mut1: GAA to CAA) had no effect on exon 9 exclusion (Figure 2B). In contrast, more severe disruption of the 5’-GGGGAAU-3’ motif with a change of Gly-Glu to Ala-Asp (mut3: GGGGAA to GCGCAU) almost abolished aberrant splicing of exon 9. A change of the motif to Arg-Val (mut4: GGGGAA to CGCGUA) completely abolished aberrant exon 9 splicing and had the same effect as a complete deletion (mut7) of the ESS motif (Figure 2B). Since these mutations do not affect the UG-rich TDP-43 binding site in intron 8 and since TDP-43 binding is indispensible for aberrant splicing, we conclude that the 5’-GGGGAAU-3’ motif constitutes a novel ESS in exon 9 which displays a sequence with a strong potential for hnRNP A1 binding (Figure 2B).

### 3.2. Binding studies of hnRNP A1 tandem RRMs with the novel CFTR exon 9 ESS sequence

Next, we aimed to characterize the interaction between hnRNP A1 and the CFTR exon 9 ESS sequence identified by our splicing assay. We used a DNA sequence comprising the CFTR exon 9 ESS sequence and characterized the interaction using ITC measurements. The hnRNP A1 tandem RRM protein was titrated with 5’-CAGGGATTTGGGGAC-3’ ssDNA sequence (Figure 3A). The ssDNA at a concentration of 200uM was added stepwise to a 28uM protein solution. The stoichiometry represented by the N parameter is 0.736 indicating that one protein molecule binds to one molecule of ssDNA. The value of the dissociation constant (K_*d*_ = 233 nM) reveals a strong affinity between the protein and the ESS motif. We note that the reaction is spontaneous with a favorable enthalpy and entropy. The enthalpy value ΔH = -2.37e^4^ kcal/mol, is consistent with an exothermic reaction. The negative entropy value (ΔS = -56.6 kcal/mol) indicates that the reaction weighs toward the formation of a more ordered structure such as a RNA-protein complex (Figure 3A). Next, we set out to investigate the role of the individual RRMs of hnRNP A1. RRM1 domain of the hnRNP A1 protein is titrated with the 5’-CAGGGATTTGGGGAC-3’ ssDNA sequence (Figure 3B). The protein concentration in the cell is 25uM and the ssDNA concentration is 200uM. The N value of 0.496 indicative of two protein molecules binding to one ssDNA which is in line with two binding sites in the ESS motif. The K_*d*_ of 5.8μM reveals a weaker affinity as compared to the tandem RRMs. The RRM2 domain of the hnRNP A1 protein was also titrated with 5’-CAGGGATTTGGGGAC-3’ ssDNA sequence (Figure 3C). The protein (25 μM) is in the cell and the ssDNA (200 μM) is again titrated to the protein solution. The N value of 0.452 also points toward two protein molecules binding to the two ESS binding sites. The K_*d*_ = 6.8μM also indicates a weak affinity. The fact that both isolated RRMs bind the ESS sequence with micromolar affinity as compared to the tandem RRMs which bind in the submicromolar range, is consistent with a cooperative binding mode of the tandem RRMs previously described for other ESS sites (Figures 3A-C)^30^.

**Figure 3.**
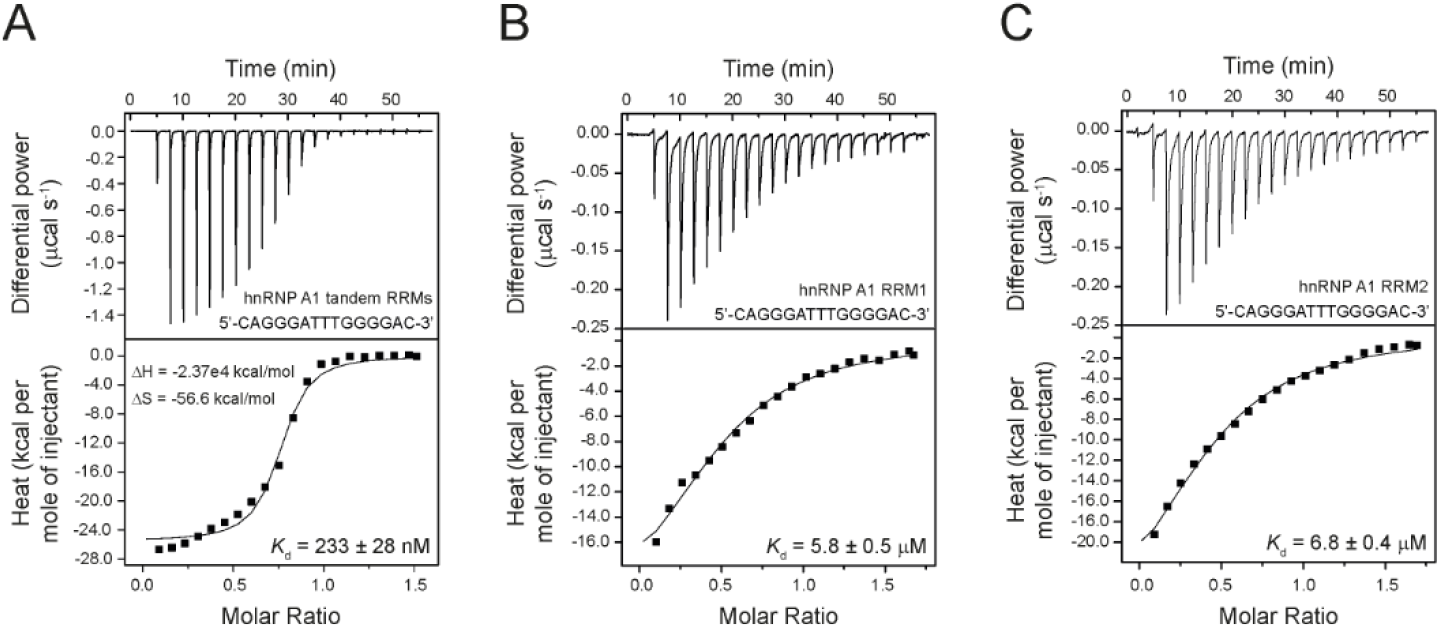
hnRNP A1 binds the novel bipartite ESS motif in CFTR exon 9. (**A**) Affinity of the hnRNP A1 tandem RRMs for the bipartite ESS motif was determined by ITC. The determined *K*_d_ value is indicated with standard deviation as well as ΔH and ΔS. (**B**) Affinity of hnRNP A1 RRM1 for the bipartite ESS motif was determined by ITC. Two copies of hnRNP A1 RRM1 bind to the bipartite ESS motif. The determined *K*_d_ value is indicated with standard deviation. (**C**) Affinity of hnRNP A1 RRM2 for the bipartite ESS motif was determined by ITC. Two copies of hnRNP A1 RRM2 bind to the bipartite ESS motif. The determined *K*_d_ value is indicated with standard deviation.

### 3.3. Binding studies of the individual hnRNP A1 RRMs with the separated ESS motifs

To further investigate the interaction of hnRNP A1 with the ESS sequence, we divided the ssDNA into two segments. We measured the affinities for both parts of the ESS motif (5’-CAGGGAT-3’ and 5’-TGGGGAAT-3’). The ITC experiments shows that the binding of RRM1 for 5’-CAGGGAT-3’ and 5’-TGGGGAAT-3’ ssDNA occurs at a 1:1 ratio with a *K*_d_ values of 0.85 μM and 7.25μM, respectively (Figures 4A-B).

**Figure 4.**
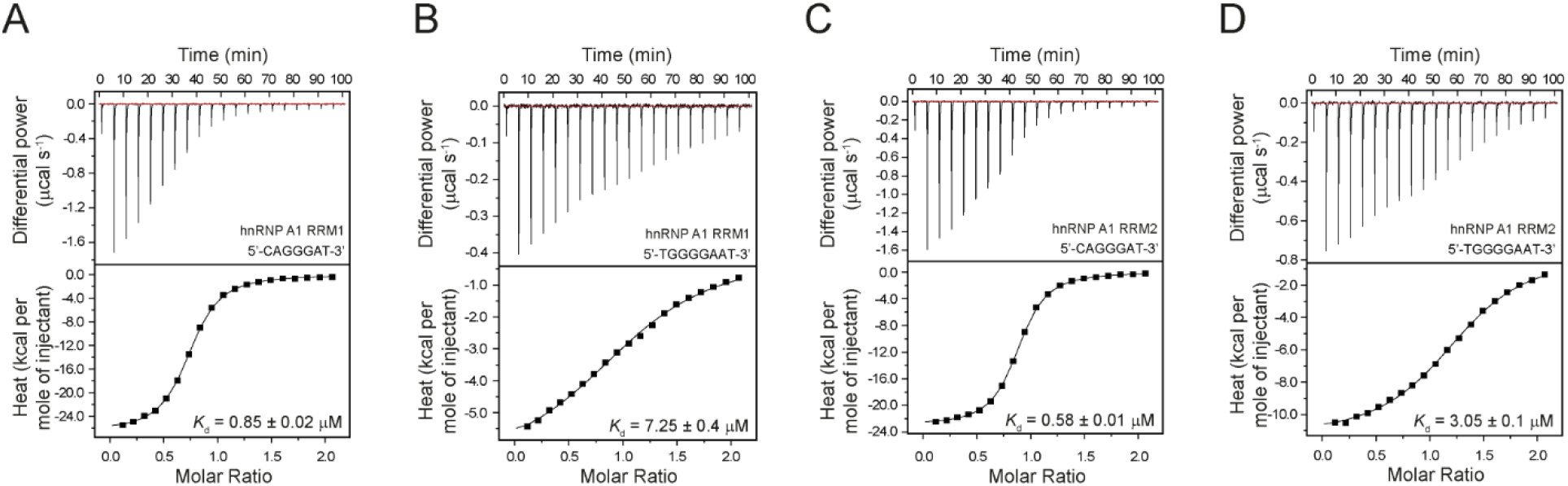
Both hnRNP A1 RRM1 and RRM2 bind the individual segments of the ESS motif of CFTR exon 9. (**A-B**) Affinity of hnRNP A1 RRM1 for the individual segments of the ESS motif were determined by ITC. 1:1 complexes are formed with the individual segments of the ESS motif. The determined *K*_d_ values are indicated with standard deviation. (**C-D**) Affinity of hnRNP A1 RRM2 for the individual segments of the ESS motif were determined by ITC. 1:1 complexes are formed with the individual segments of the ESS motif. The determined *K*_d_ values are indicated with standard deviation.

Thus, the RRM1 domain has an almost ten-fold higher affinity for the first part of the bipartite ESS motif. For RRM2, the ITC data show that the binding of 5’-CAGGGAT-3’ and 5’-TGGGGAAT-3’ DNA occurs also at a 1:1 ratio with a *K*_d_ value of 0.58 μM and 3.05 μM, respectively (Figures 4C-D). In conclusion, the RRM2 domain has a five-fold higher affinity for the first part of the ESS motif as well. Nonetheless, the similar binding affinities of the individual RRM domains for the separated ESS motifs do not allow to reach a conclusion about the orientation of the ESS motif upon interaction with the tandem RRM scaffold of the hnRNP A1 protein.

### 3.4. Assessment of the ESS – hnRNP A1 interaction by solution NMR spectroscopy

We used the assignment of the hnRNP A1 protein published by the Allain group and superimposed them onto our ^1^H-^15^N HSQC spectra of the amide fingerprint region of hnRNP A1^22^. We produced four complexes comprising each separated RRM domain with each of the separated ESS segments.

The ^1^H-^15^N HSQC spectra of the four complexes were recorded at 298K. Since the second ESS segment contains four guanines which could form a G-quadruplex irrelevant to exon 9 splicing, and since previous studies showed that G-quadruplexes bind to the hnRNP A1 protein with high affinity^*31*^, we mutated the second ESS motif from 5’-TGGGGAAT-3’ to 5’-CTGGGCAC-3’ according to the mut1 which had no effect on aberrant splicing (Figure 2B).

With the titration results, we could confirm the binding of the RRM1 domain with the 8-mer ssDNA model sequence (5’-CAGGGATC-3’) (Figures 5A-B). The ssDNA binding of the free protein shown in blue, can be followed upon stepwise addition of the ssDNA (Figure 5A). The chemical shift perturbations (CSP) observed at each titration point indicate the potential ssDNA binding site on the protein and that the interaction occurs in a fast exchange regime in the NMR timescale. We considered all the CSPs > 0.06 more than average, indicating that the affected amino acids are either involved in the interaction with the 8-mer ssDNA model or indirectly affected by ssDNA binding. The affected amino-acids are Gln13, Leu17, Gly20, Phe24, Glu29, Ser30, Thr42, Val45, Val46, Thr52, Ser55, Gly57, Ala72, Asn74, His78, Lys88, Ala90, Val91, Ser92, and Arg98 (Figure 5B). Mapping these CSPs onto the structure shows that the interaction site of the ssDNA mostly comprises the beta sheet surface of the protein which is the canonical RNA interaction site on the RRM fold.

**Figure 5.**
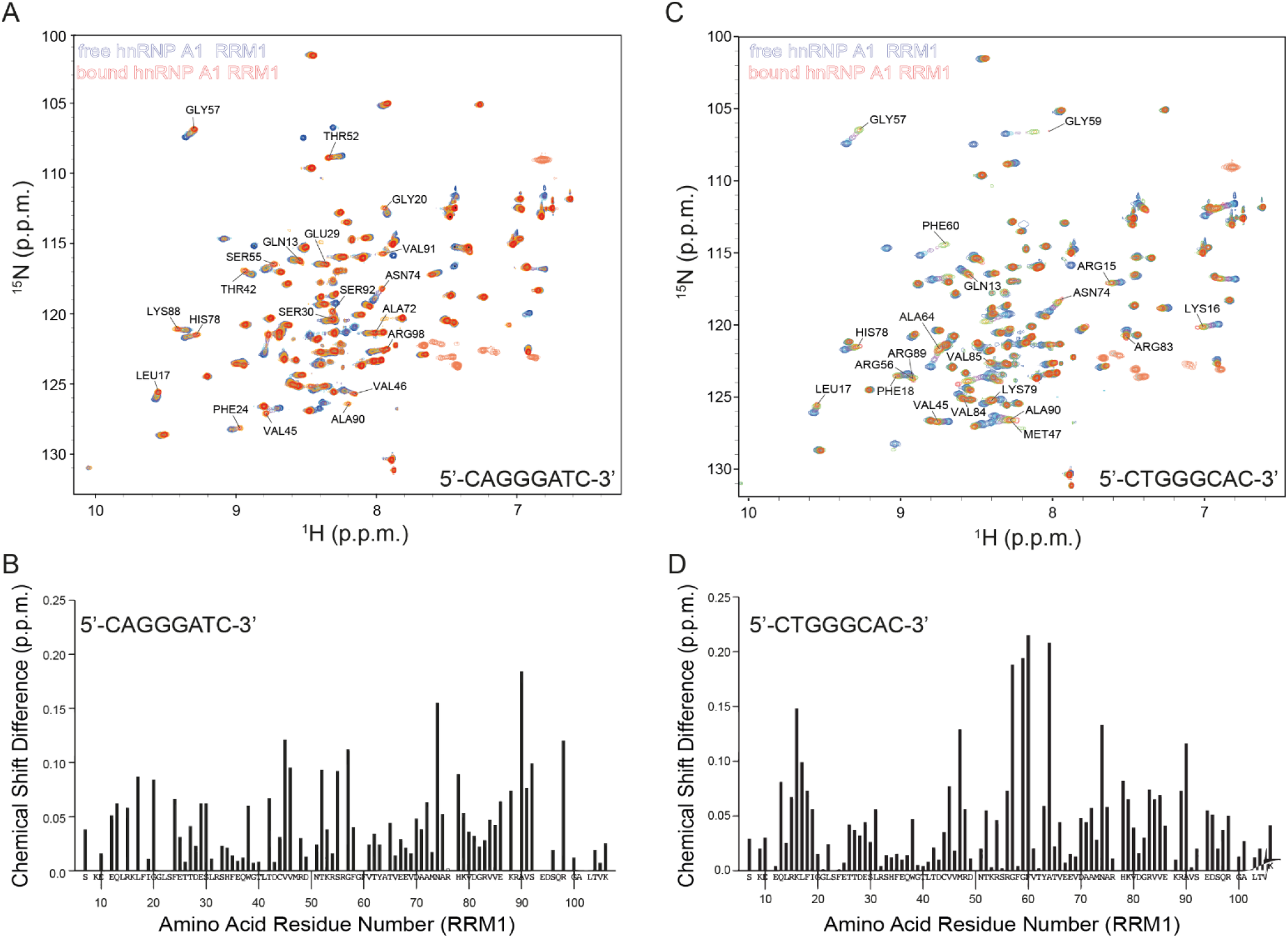
NMR titration of hnRNP A1 RRM1 reveals specific interactions with the individual segments of the ESS motif. (**A** and **C**) Superimposition of _1_H-_15_N HSQC spectra of _15_N-labelled hnRNP A1 RRM1 in the free (blue) and short ESS motif-bound form (red). The amide cross-peaks shifted upon interaction are labelled with amino acid type and residue number. (**B** and **D**) Combined chemical shift perturbations Δδ=[(δH_N_)_2_ + (δN/6.51)_2_]_1/2_ of hnRNP A1 RRM1 amide resonances upon short ESS motif binding versus the hnRNP A1 RRM1 amino acid sequence.

We subsequently recorded ^1^H-^15^N HSQC spectra (298K) to monitor the binding of the RRM1 domain with the 8-mer DNA model (5’-CTGGGCAC-3’) corresponding to the second part of the ESS motif (Figure 5C). We used the same ssDNA titration steps as for the first ESS motif to monitor ssDNA binding. Again, the nature of the CSPs indicated a fast exchange regime at the NMR time scale that the RRM1 domain binds also to the second part of the ESS motif. All the CSP > 0.06 are considered more than average and involve amino-acids Gln13, Arg15, Lys16, Leu17, Phe18, Val45, Met47, Arg56, Gly57, Gly59, Phe60, Ala64, Asn74, His78, Lys79, Arg83, Val84, Val85, Arg89, Ala90 (Figure 5D). The interaction site of the ssDNA comprises mostly the beta sheet of the protein which are again the canonical RNA interaction sites on the RRM fold.

Next, we monitored binding of RRM2 to the separate ESS motifs. Interaction occurs in a fast exchange regime in the NMR time scale. The NMR titration confirmed the interaction between RRM2 and the first ESS motif (5’-CAGGGATC-3’) as well as the second ESS motif (5’-CTGGGCAC-3’) (Figures 6A and 6C). Thus, our NMR titration experiments confirm the ITC results which showed that both individual RRMs interact with each part of the ESS motif. The CSPs > 0.05 observed upon binding of RRM2 to the 8-mer ssDNA (5’-CAGGGATC-3’) involve amino acids Leu103, Thr104, Lys106, Lys107, Ile108, Val110, Lys131, Glu136, Thr139, Asp140, Gly142, Gly144, Phe151, Ser159, Asp161, His169, Lys180, Gln185, Met187, Ser189 (Figure 6B).

**Figure 6.**
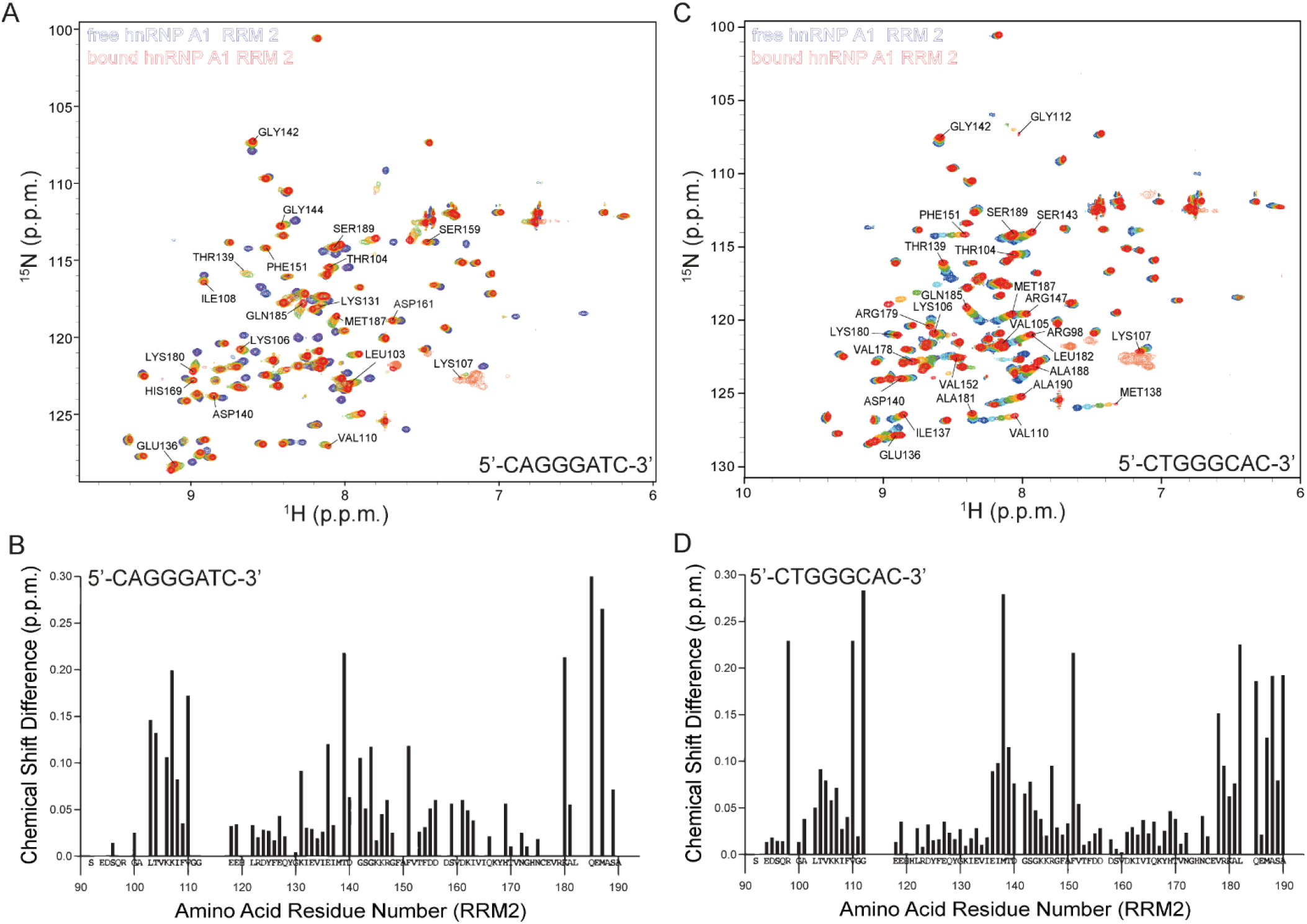
NMR titration of hnRNP A1 RRM2 reveals specific interactions with the individual segments of the ESS motif. (**A** and **C**) Superimposition of _1_H-_15_N HSQC spectra of _15_N-labelled hnRNP A1 RRM2 in the free (blue) and short ESS motif-bound form (red). The amide cross-peaks shifted upon interaction are labelled with amino acid type and residue number. (**B** and **D**) Combined chemical shift perturbations Δδ=[(δH_N_)_2_ + (δN/6.51)_2_]_1/2_ of hnRNP A1 RRM2 amide resonances upon short ESS motif binding versus the hnRNP A1 RRM2 amino acid sequence.

Likewise, the CSPs > 0.05 observed upon binding of RRM2 and the 8-mer ssDNA (5’-CTGGGCAC-3’) involve a similar region on the protein (Arg98, Thr104, Val105, Lys106, Lys107, Val110, Gly112, Glu136, Ile137, Met138, Thr139, Asp140, Gly142, Ser143, Arg147, Phe151, Val152, Val178, Arg179, Lys180, Ala181, Leu182, Gln185, Met187, Ala188, Ser189, Ala190 (Figure 6D). The interaction sites of both ssDNAs comprise mostly the beta sheet of the protein as well as the C-terminal helix.

Next, we used the CSP results from the NMR titration experiments of the individual RRMs and the separated ESS segments to visualize the potential ESS binding site on the hnRNP A1 tandem RRMs structure determined by the Allain group^22^. The overall fold and arrangement of the tandem hnRNP A1 RRMs presents a continuous binding site for the novel, bipartite CFTR exon 9 ESS segment (Figure 7). Despite that fact that our experiments did not reveal a preference of one of the RRMs for a certain segment of the bipartite ESS sequence, both possible orientations of the ESS segment on the tandem RRMs cover the same continuous protein surface and could thereby impair access of canonical splicing factors to the CFTR exon 3’ss and subsequently cause aberrant CFTR exon 9 splicing.

**Figure 7.**
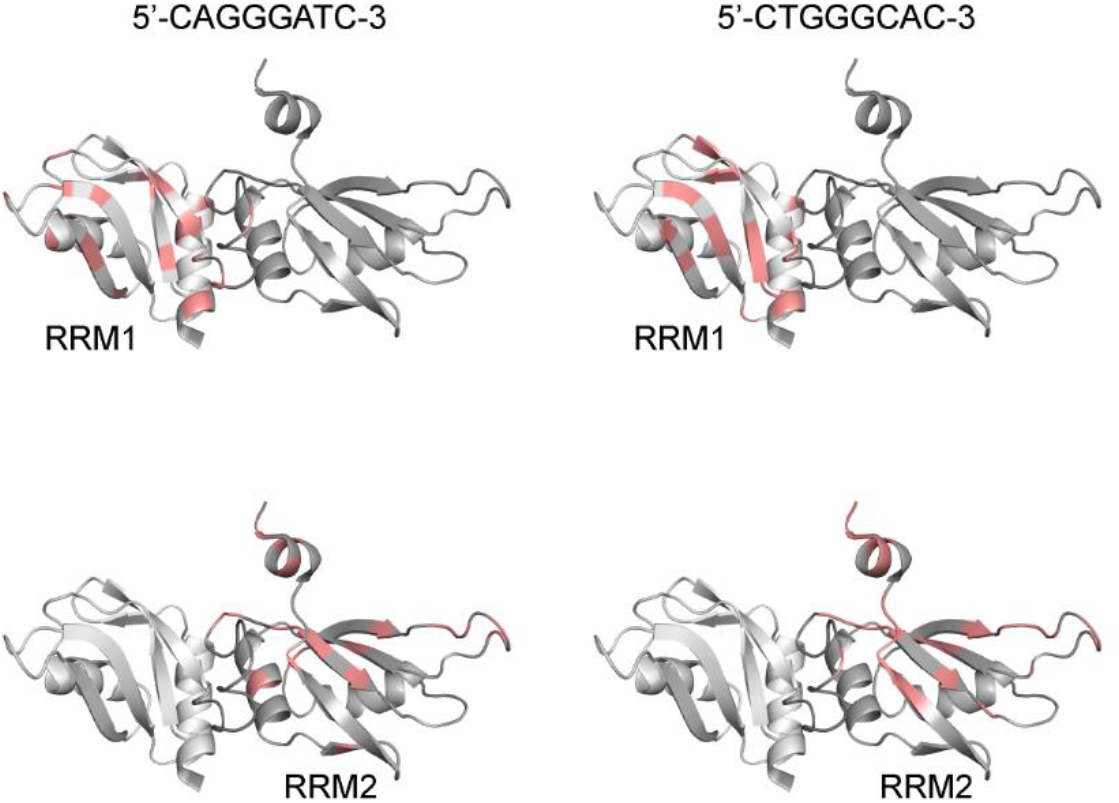
The hnRNP A1 tandem RRMs display a continuous binding site for the novel CFTR exon 9 ESS motif. The CSPs from the titration experiments of the individual hnRNP A1 RRMs and the short segments of the ESS motif are displayed onto the structure of the hnRNP A1 tandem RRMs (PDB ID: 2lyv). RRM1 is coloured in light gray and RRM2 in dark gray. The CSPs are indicated onto the ribbon structure in salmon and the sequence of the corresponding ESS segments is displayed on top.

## 4. Discussion

Alternative splicing is an important corner stone of RNA-based, posttranscriptional regulation of gene expression. Production of diverse protein products through inclusion and exclusion of certain exons from the same pre-mRNA is a major contributor to the complexity of higher organisms. But at the same time, splicing also depends on correct RNA and protein products for proper functioning and thus variations in both components can lead to missplicing and disease^32^. One such variant is found in the polymorphic (TG)_m_(T)_n_ locus upstream of CFTR exon 9 within intron 8 which leads to recruitment of TDP-43 to the pre-mRNA and subsequent TDP-43-dependent recruitment hnRNP A1. The UG-rich binding site of TDP-43 in the CFTR pre-mRNA is well characterized both functionally^15, 29^ and structurally^16-17^ and it is known that hnRNP A1 is recruited to the 3’ss *via* a protein-protein interactions with TDP-43^18-19^. The molecular origin of CFTR exon 9 splicing inhibition *via* TDP-43-dependent recruitment of hnRNP A1, on the other hand, remained elusive.

Our study identifies a new ESS comprising a bipartite sequence motif at the CFTR exon 9 3’ss and the 5’end of exon 9 (5’**-**<u>AG</u>GGAUUUGGGGAAU-3’). We show that hnRNP A1 is able to specifically interact with this bipartite ESS motif and that the integrity of this site is instrumental for aberrant splicing of CFTR exon 9. Our minigene splicing assay shows that the wt sequence around the CFTR 3’ss produces several bands corresponding to the inclusion and skipping of exon 9. Four different mutations in exon 9 which alter the 5’-GGGGA-3’ motif but leave the 3’ss intact, clearly show that variation of this sequence affects the inclusion of exon 9 in the mRNA. Mutating the 5’-GGGGA-3’ motif to 5’-CGCGT-3’ as well as its complete deletion are associated with non-skipping of exon 9, i.e. the full inclusion into the final mRNA. Thus, these experiments reveal for the first time an ESS motif in CFTR exon 9 causative for exon 9 skipping.

We also investigated the RNA-binding mode of hnRNP A1 to the ESS that silences splicing of CFTR exon 9. The ITC and NMR titration experiments strongly suggest that both RRMs of hnRNP A1 interact with the bipartite ESS motif. The affinities obtained from ITC experiments using the isolated RRMs and single binding sites of the bipartite ESS motif are comparable, suggesting that there is no preferred sequence motif for the individual RRMs determining the orientation of hnRNP A1 on the pre-mRNA target. Previous studies have shown that the hnRNP A1 protein can bind specifically and with high affinity to ssRNA or ssDNA sequences because there are no qualitative differences between them^21^ and several high-affinity sequences have also been discovered for hnRNP A1. Indeed, SELEX experiments isolated a high-affinity RNA sequence of 5’-UAGGGA/U-3’ that is recognized by hnRNP A1^27^ which is similar to the CFTR exon 9 ESS motif. The specific recognition of the motif was confirmed by an x-ray crystallography structure of hnRNP A1 tandem RRMs bound to a telomeric ssDNA sequence (5’-TTAGGGTTAGGG-3’)^21^. The structure revealed that both RRM1 and RRM2 of the same monomer interact with two different strands of ssDNA, in an antiparallel manner. In this structure, RRM1 binds to four nucleotides (5’-TAGG-3’), while RRM2 preferably binds to five nucleotides (5’-TTAGG-3’). The CFTR exon 9 ESS motif also contains two binding sites with similar sequence but our NMR and ITC data suggest that a single copy of hnRNP A1 binds both motifs *via* the tandem RRMs. Thus, the dimerization in the x-ray structure might be just caused by crystal packing forces and not be relevant for CFTR exon 9 skipping.

A study performed by Allain and colleagues has identified optimal recognition motifs that bind the individual RRMs^33^. RRM1 binds to the sequence 5’-U/CAGG-3’ and RRM2 to the sequence 5’-U/CAGN-3’. They also show that both RRMs of hnRNP A1 bind to their RNA target containing the two AG-binding sites without dimerization of the protein and that hnRNP A1 binds the ISS-N1 pre-mRNA with directionality where RRM2 interacts with the 5’ motif and RRM1 binds the 3’ motif.

Another study investigated the interaction between hnRNP A1 to an RNA target, namely the pri-mir-18a. Again, this interaction involves both RRMs and a region encompassing two 5’-UAG-3’ motifs^34^. In this study, it was demonstrated that cooperative binding of both domains to the related RNA motifs results in a strong enhancement of binding affinity and allows the unwinding of the RNA target as a stem loop. Our ITC experiments also reveal that individual RRMs interact with the ESS in the low micromolar range while the tandem RRMs display nanomolar affinity consistent with cooperative binding of both RRMs to the bipartite ESS.

Further work is needed to determine the orientation of hnRNP A1 along the CFTR exon 9 ESS element. Binding this natural target with directionality might require the protein-protein interaction of full-length TDP-43 and hnRNP A1 mediated by their glycine-rich C-terminal domains^18-19^. Consistent with this idea, disrupting this interaction *via* mutations or deletion in the C-termini of TDP-43 as well as hnRNP A1, abolishes exon 9 skipping despite the fact that hnRNP A1 maintains its ability to bind to the target CFTR exon 9 ESS element. Studying the RNA-protein and protein-protein interactions within the entire aberrant splice site using full-length TDP-43 and hnRNP A1 is required to shed light on the precise mechanism of aberrant splicing occurring at CFTR exon 9.

## Author Contributions

P.J.L. designed the project. C.B. and M.Y.C. expressed and purified proteins and purified ssDNA. C.B. performed all NMR measurements, ITC experiments with 15-mer ssDNA and analyses. M.Y.C. performed all ITC experiments with individual RRMs and shorter ssDNAs. C.S. performed all the minigene splicing assays and analyzed the data. C.B. wrote an initial draft, P.J.L. wrote the manuscript and made proper figures, E.B. and C.S. wrote the minigene splicing assay part and made corrections to the text. All authors discussed the results and approved the manuscript.

## Data Availability Statement

The authors confirm that the data supporting the findings of this study are available within the article.

## Acknowledgments

We wish to thank Prof. Radova Fiala from Josef Dadok National NMR Centre for the skillful NMR technical assistance. We acknowledge CF BIC and CF NMR, Instruct-CZ Centre, supported by MEYS CR (LM2018127) and European Regional Development Fund-Project „UP CIISB” (No. CZ.02.1.01/0.0/0.0/18_046/0015974).

